# Eye tracking impairments in children with protein-energy malnutrition

**DOI:** 10.1101/2021.01.08.425941

**Authors:** Natalia L. Almeida, Jessica B. S. Silva, Nayara P. Silva, Thiago P. Fernandes, Natanael A. Santos

## Abstract

Background: Malnutrition is characterized by impaired nutrient ingestion and absorption, and it is still one of the most substantial causes of morbidity and mortality in children worldwide. To the best of our knowledge, this is the first study investigating eye tracking in children with protein-energy malnutrition (PEM). We sought to investigate how PEM may affect eye movement. Methods: Twenty children without PEM (mean age = 10.8; SD = 1.0 years) and 18 children with PEM (mean age = 10.9; SD = 1.2 years). Here we used three types of tests or stimuli: one formed by a maze and two by seven errors games (boats and elephants). Results: Our results indicated that children with PEM had impaired performance on all of the tests used here. These data suggest that the nutritional impairments during the first year of life, the critical period in visual system development, can have direct impact on eye movement patterns. Conclusions: Our findings must be replicated so that neurophysiological patterns of PEM can be precisely understood. However, this study has repercussions in several areas of knowledge.

## 1. Introduction

Nutrition is fundamental during the critical period of development due to the rapid corporal growth, serving as an indicator of growth and development [1, 2]. Malnutrition, in turn, refers to a condition caused by the loss in nutrient ingestion or absorption, such as proteins, carbohydrates, fats, vitamins and mineral salts [3, 4]. Since the organism needs nutrients to support development, both the brain and the nervous system are vulnerable to the effects of malnutrition [3].

Studies indicate that malnutrition may affect brain plasticity, affecting cognitive functions such as memory, executive functions and thought [5, 6]; likewise, malnutrition may also affect visual processing [7]. Some studies point, for instance, that the loss of amino acids and its precursors in the organism, mainly of glutamate, GABA and dopamine, may affect neurotransmission [8].

Neurophysiological changes caused by malnutrition also happen in the visual system. For instance, we may mention: (1) changes to the lens and retinae, (2) reduction in size and number of optical fibres (3) reduction in number of synapses per neuron and in total number of synapses in the visual cortex [9]. It is important to highlight that vision develops mainly in the first years of live and its sensorial plasticity is greater in the first two years. Until this age, any obstacle to vision development may cause late changes [10]. In this way, systematic and objective measures to evaluate the existence of progressive visual changes are important, such as sensitivity to contrast, or measures such as eye tracking.

Eye tracking is a technique in which the eye movements may be monitored during the execution of several tasks. Some types of movement may be described and identified, however the most frequent and used in eye movement registering systems are those called fixations and saccadic movements [11]. Saccadic movements are very fast and voluntary (approximately two to four seconds between fixations) and serve to direct the fovea to the visual processing zone of interest. Fixations are quick and momentaneous stops of the fovea (approximately 100 ms) in a specific area of the visual scene to capture and process information. The time of fixation may indicate the accuracy and quantity of visual processing, which may be used to investigate congenital or acquired problems. Research suggests that the decision points are generally followed by long periods of fixations, when the cognitive processes nullify visual perception [12].

Few studies with psychophysical methodology have given due attention to the impact of protein-energy malnutrition (PEM) in visual functions [13, 14]. Given that malnutrition is still one of the substantial causes of morbidity and mortality of children from all over the world, and considering the paucity in studies involving eye tracking in children afflicted with PEM, it is important to investigate new aspects of this condition. In this sense, we sought to investigate the possible effects of early malnutrition, more specifically PEM, in eye tracking using different tests. We expect that children with PEM should present lower performance in the three tests used here, when to healthy children.

## 2. Method

### 2.1 Participants

Thirty-eight children participated in this study, 18 with prior PEM and 20 healthy controls, some former classmates of the PEM group, who had no histories of malnutrition, aged between 7 and 12 years old. Children from both groups were recruited from public school and had normal or corrected-to-normal vision as determined by visual acuity of at least 20/20 [15].

In the Study group, children within the cut-off value for PEM according to the WHO parameters were included [16]. We also employed Waterlow’s criteria to investigate agreement between parameters [17]. The use of these parameters demands the use of reference standards and cut-off value based in the following indices: height/age (obtained by comparison, according to gender, between the observed height and the one expected for the chronological age) and weight/height (obtained, comparatively, according to gender, weight and expected height for age). In the control group, children with similar characteristics to those of the study group (age, gender, same school year and public education), except for nutritional condition, were included.

Children with records of or suspected ocular, neurological, cardiovascular diseases, diabetes, hepatitis or an identifiable neuropsychological condition at the time of screening were excluded. In addition, children with birth weight < 2500g, presence of pre- or post-natal complications or other serious diseases after the first year of life were excluded.

This research followed the ethical principles of the Declaration of Helsinki and the Committee of Ethics in Research of the Health Sciences Centre of Federal University of Paraiba (CAAE: 14798613.5.0000.5188). Written informed consent was obtained from all of the participants (Assent Term for subjects younger than 18 years old, duly signed by parents or guardians.

### 2.2 Instruments

#### 2.2.1 Eye tracking - Eyetracker Tobii TX300

A 300 Hz eyetracker was used, measured binocularly. It was integrated to a 23’’ monitor (maximum resolution of 1920 × 1080 pixels and 300 cd/m^2^) where stimuli were presented for the subject to visualise. This equipment was connected to a Dell Latitude 3450 notebook with a 14.0” HD (1366 × 768) and Windows 8.1 Pro 64 bits operational system, Intel® Core™ i7 -(5500U 2.4 GHz, 8 GB of RAM memory) used for visualisation by the researcher. The *Tobii Studio* software was used for test elaboration, recording and eye movement analysis.

At the start of eye calibration, the equipment measures the distance between the participant’s eyes and the screen where the stimuli are presented. In this study, the measure used was 65 cm (as indicated by the instrument’s manual). Calibration occurs through the fixed gaze in a red ball that moves around the screen. After that, the equipment begins calibration, which serves to adjust measures to the geometric characteristics of the eyes of each participant. If there are points in the screen not adequately calibrated to the real position of the eyes, the equipment indicates these points with green lines, and the researcher may repeat their calibration. Once all points are calibrated correctly (with no green lines), the calibration configurations are saved and the test begins.

### 2.3 Stimuli

#### 2.3.1 Maze test

This test seeks to evaluate the eye movement and decision making [12]. The test presents regions that, even in a short time period, let one decide which path to take (decision point). The beginning of the test (point A) coincides with the centre of the screen and of the maze, initial fixation point, the standard adopted by the eye tracker. It possesses four arrival points (B points) with a symmetric degree of difficulty. There are two different types of path in the maze, with the same degree of difficulty (for more information on the maze, see Santos et al., [12]. The stimulus used in the test may be visualised in Fig. 1.

**Fig. 1.**
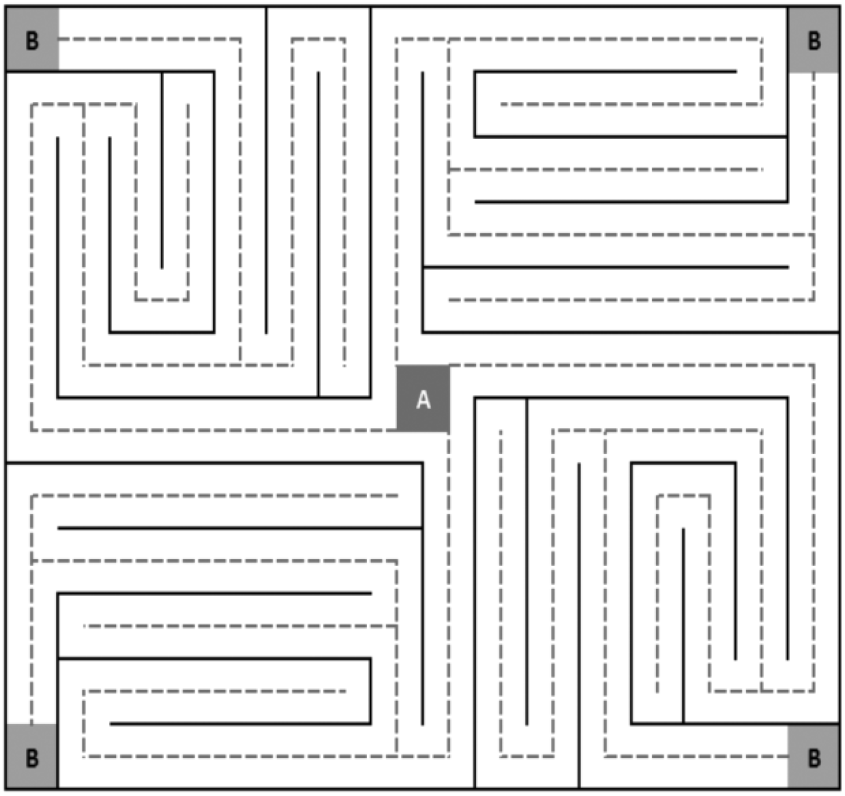
Example of stimulus used in the maze test.

#### 2.3.2 Seven errors test

Also adapted by our lab, it seeks to evaluate eye movement. The stimuli are relatively simple and were planned to monitor eye movement, specifically the fixations and saccades. The task is composed by two different pairs of images: a pair referring to an image of a boat (one being the original and the other containing the image with the seven errors, see Fig. 2A), and the other referring to the image of an elephant (one being the original and the other containing the image with the seven errors, see Fig. 2B). The original image and the image containing the seven errors were presented simultaneously in the screen, side by side so that participants could compare them and identify the errors. The stimuli were selected based on the standardized set proposed by [18].

**Fig 2.**
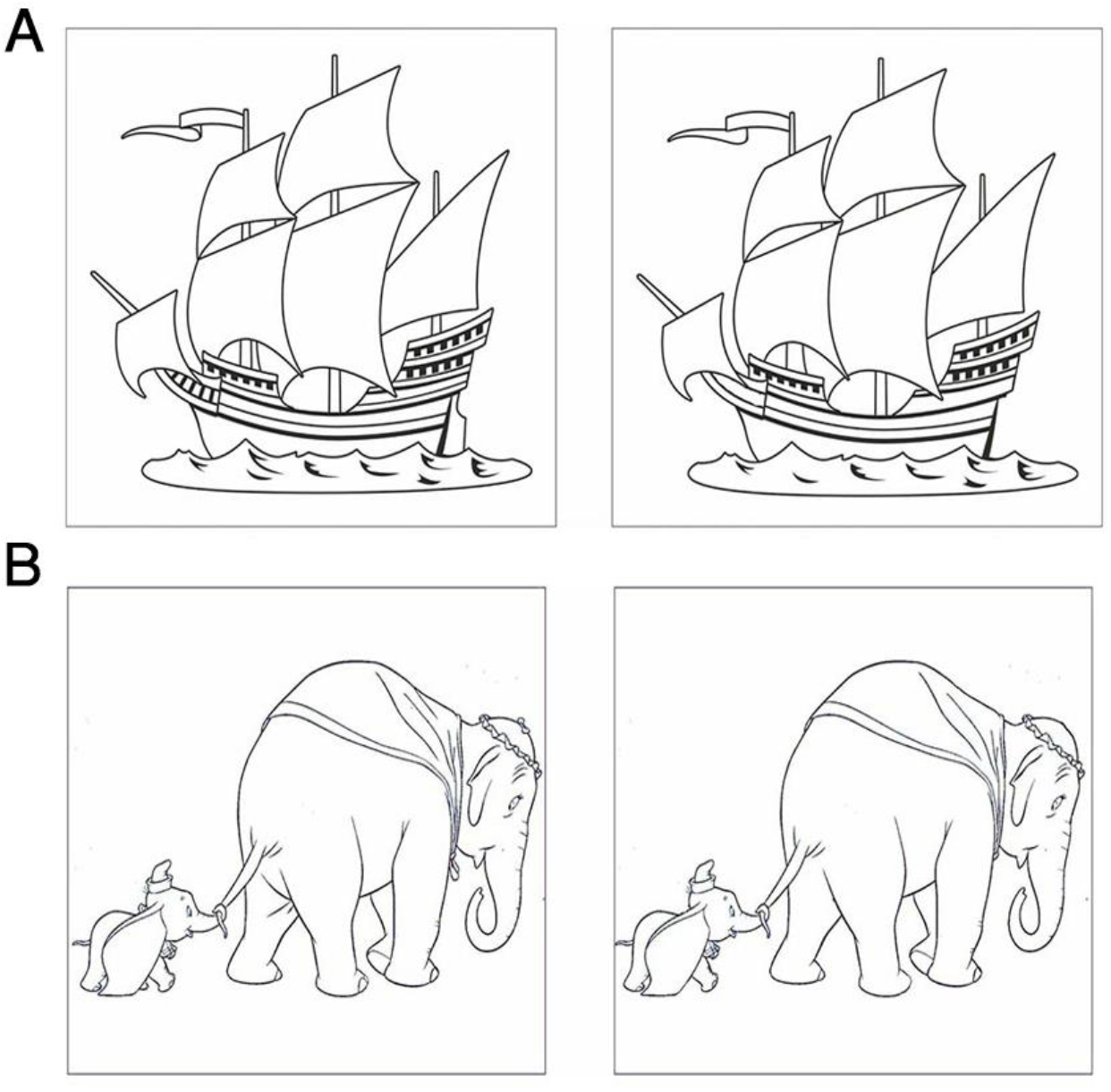
Examples of stimuli used in the seven errors test, the boat (A) and the elephant (B).

### 2.4 Procedures

The recruitment of the children was done in public schools through the director board, which mediated the contact of the researching team with the guardians. The guardians who authorised the children’s participation in the study signed the terms and the children participating in the study were subject to the anthropometric evaluation.

The children were instructed to sit 65 cm from the monitor in a comfortable, but steady position, and to not make abrupt movements with their heads. Initially, the system calibration was made in order to verify correspondence between the eye position as calculated by the eye tracker and the actual eye position. After calibration, the instructions were exhibited in the screen.

After understanding the instructions, the participants were instructed to gaze in a central point of the screen, and, immediately afterwards, to the stimulus. Concerning the maze test, the participants were asked to go from point A to point B. Concerning the seven errors test, as the first pair of images was presented, the participant was tasked with finding the seven errors in the stimulus. The execution time of the maze test depended on arrival to point B. Concerning the seven errors test, the first part ended when all errors were found or when the maximum time of two minutes had elapsed. The stimulus presentation order was random.

### 2.5 Data analysis

For each condition, data distribution was assessed using measures of central tendency and measures of dispersion. Distributions for each group were compared using the Monte Carlo method for skewness and kurtosis, and the cutoff value was > 1.96 [19–21]. Statistical analysis was performed using SPSS 25.0.

Both groups’ data were characterized by normal distributions, and so parametric tests were performed to analyse continuous data. To compare groups on the nominal variable gender the non-parametric chi-squared test was conducted. Three separate multivariate analyses of variance (MANOVA) were conducted to (i) analyse the results of the maze test (three dependent variables), (ii) analyse the test results of the seven errors test for the elephant stimuli (three dependent variables), and (iii) analyse the test results of the seven errors test for the elephant stimuli (three dependent variables). Omega squared (ω^2^) was used to assess effect sizes (for small sample sizes, ω^2^ reduces bias; [22]. The estimate of effect size for pairwise analyses occurred using the Hedges’ *g* (indicated for smaller samples). Bonferroni's correction was applied to adjust the *p*-value and reduce the risk of Type 1 errors. Using the *Bonferroni* correction to adjust the *p* value and avoiding type I errors. Correlation analyses were made, separately, in order to investigate association between the body mass index and total time (in seconds) to complete the tests in the malnourished children group.

## 3. Results

### 3.1. Biosociodemographic data

The groups did not differ in age, [*t* (36) = .240, *p =* .82] or level of education, [*t* (36) = .239, *p =* .82], or gender [*χ* (1) = .145, *p =* .71]. The sample had a majority of female participants, with mean age of 10.9 years. The body mass index (BMI) had mean score of 19.17. The table below shows the data average in each group (Table 1).

**Table 1.** Biosociodemographic data of the sample and outcomes of the tests.

| Data distribution | Control group ($n = 20$) | Study group ($n = 18$) |
| --- | --- | --- |
| **Biosociodemographic data** |  |  |
| Gender (%) |  |  |
| Boys | 9 (45%) | 7 (35%) |
| Girls | 11 (55%) | 11 (65%) |
| Age (SD) | 10.80 (1.01) | 10.89 (1.27) |
| Height, cm (SD) | 152.20 (0.06) | 143.61 (0.09) |
| Weight, kg (SD) | 50.93 (7.33) | 33.63 (6.43) |
| BMI (SD) | 22.14 (2.84) | 15.87 (0.92) |
| **Maze test** |  |  |
| *Amount of fixation* |  |  |
| Mean (SD) | 27.00 (8.16) | 59.44 (21.27) |
| Skewness | 0.20 | 0.47 |
| Kurtosis | 0.65 | 0.47 |
| *Average fixation duration (ms)* |  |  |
| Average (DP) | 293.33 (67.74) | 353.60 (71.33) |
| *Skewness* | -0.14 | 0.30 |
| *Kurtosis* | -0.27 | -0.65 |
| *Total spent (s)* |  |  |
| *Average (DP)* | 9.65 (4.32) | 22.14 (8.44) |
| *Skewness* | 1.19 | 1.14 |
| *Kurtosis* | 0.64 | 0.36 |
| **Seven errors test** |  |  |
| *Amount of fixation – Boat / Elephant* |  |  |
| *Average (DP)* | 164.55 (80.84) / 192.90 (75.09) | 420.28 (92.82) / 270.61 (75.82) |
| *Skewness* | 1.22 / 0.64 | 0.55 / 0.42 |
| *Kurtosis* | 0.45 / 0.28 | 1.04 / 0.50 |
| *Average fixation duration (ms) – Boat / Elephant* |  |  |
| *Average (DP)* | 290.95 (43.31) / 292.50 (48.04) | 410.72 (40.42) / 382.28 (130.67) |
| *Skewness* | 0.67 / 0.98 | 0.45 / 1.01 |
| *Kurtosis* | 0.78 / 0.95 | 1.04 / 0.84 |
| *Total spent (s) – Boat / Elephant* |  |  |
| *Average (DP)* | 50.53 (17.25) / 46.60 (15.59) | 145.05 (31.26) / 78.82 (35.04) |
| *Skewness* | 0.12 / 0.46 | 0.84 / 0.73 |
| *Kurtosis* | 0.67 / 0.52 | 0.85 / 1.84 |

Note: SD = standard deviation; ms = milliseconds; s = seconds

### 3.2 Maze test

There were significant differences for all the three dependent variables [*F* (3,34) = 50.07, *p* < .001, Pillai’s Trace = .64, ω^2^ = .794; 95% CI: .688 – .856]. Discriminant analysis showed that the study group had worse performance for amount of fixation [*p* < .001, Hedges’ *g* = 1,693; 95% CI: 0.642 – 2.710], average fixation duration [*p* < .001, Hedges’ *g* = 0.853; 95% CI: 0.398 – 1.537] and total time spent [*p* < .001, Hedges’ *g* = 1.738; 95% CI: 0.679 – 2.763]. The main results are shown in Fig. 3.

**Fig. 3.**
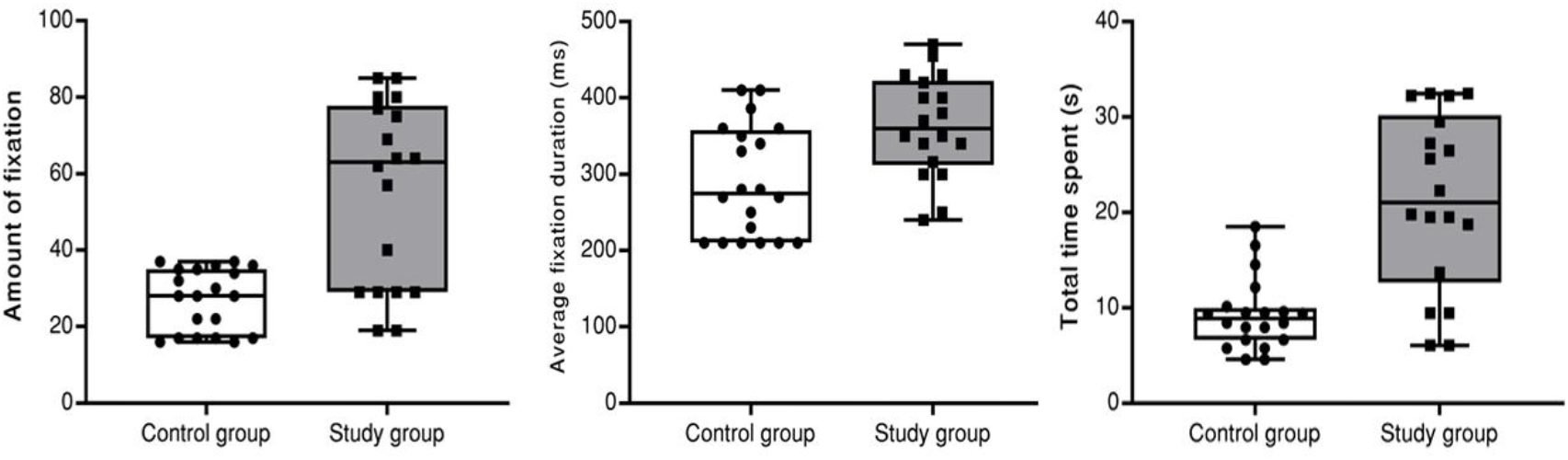
Performance of the groups for amount of fixation, average fixation duration (ms) and total time spent (s) in the maze test. The horizontal line displays the median, the boxes represent the 25-75 percentiles and the whiskers represent the range. Circles and squares represent individual participant means.

### 3.3 Seven errors test

#### 3.3.1 Boat

There were significant differences for all the three dependent variables [*F* (3,34) = 92.51, *p* < .001, Pillai’s Trace = .89, ω^2^ = .878; 95% CI: .822 – .925]. Discriminant analysis showed that the study group had worse performance for amount of fixation [*p* < .002, Hedges’ *g* = 1.656; 95% CI: 0.611 – 2.667], average fixation duration [*p* < .001, Hedges’ *g* = 2.793; 95% CI: 1.929 – 3.758], and total time spent [*p* < .001, Hedges’ *g* = 2.106; 95% CI: 0.977 – 3.199]. The main results are shown in Fig. 4A.

**Fig. 4.**
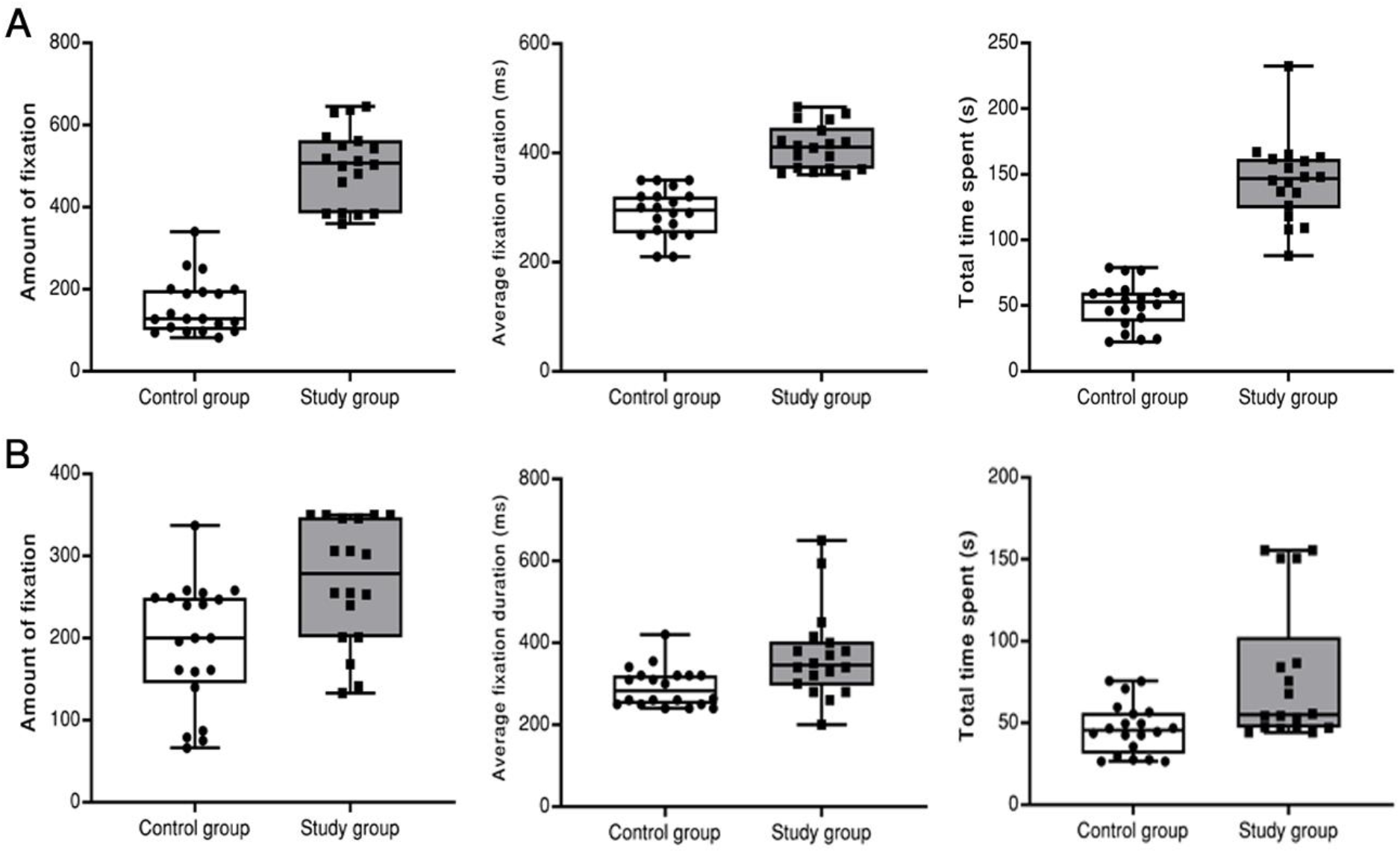
Performance of the groups for amount of fixation, average fixation duration (ms) and total time spent (s) in the seven errors test for the boat (A) and elephant (B) stimulus. The horizontal line displays the median, the boxes represent the 25-75 percentiles and the whiskers represent the range. Circles and squares represent individual participant means.

#### 3.3.2 Elephant

There were significant differences for all the three dependent variables [*F* (3,34) = 6.62, *p =* .001, Pillai’s Trace = .37, ω^2^ = .307; 95% CI: .115 – .495]. Discriminant analysis showed that the study group had worse performance for amount of fixation [*p* = .003, Hedges’ *g* = 1.277; 95% CI: 0.293 – 2.231], average fixation duration [*p* = .008, Hedges’ *g* = 1.545; 95% CI: 1.016 – 2.599], and total time spent [*p* = .003, Hedges’ g = 1.105; 95% CI: 0.454 – 2.094]. The main results are shown in Fig. 4B.

### 3.4 Correlation analyses

Results indicate correlations between the total time spent for performing the test and BMI. That is, the lower BMI, the greater the total time spent for performing the test. In other words, a lower BMI resulted in a greater time for performing the maze test (*r* = −.61, *p* < .001), the seven errors test with the elephant stimulus (*r* = −.35, *p* = .028), and seven errors with the boat stimulus (*r* = - .73, *p* < .001).

## 4. Discussion

We were able to verify for the first time, through this study, the effects of PEM on eye tracking. Our initial hypothesis that it was expected that children with early malnutrition would present a lower performance when compared to children with no PEM was supported by these findings. The results obtained with the eye tracking measurements show the existence of visual deficits in children with a record of EPM. The results for the maze test (Fig 3) and for the two seven errors tests (Fig. 4) show the same tendencies (that is, impairments to eye tracking related to previous PEM), which in a certain way support and convalidate the hypothesis as the obtained data were replicated even for a small number of children. In addition to it, the results found corroborate in a certain way other studies involving the visual system and perception indicating that children with PEM present, in general, a diminished performance when compared with healthy children. Further on, the BMI was related to this performance, where lower indices resulted in a longer time to complete the tests.

Psychophysical studies involving basic visual functions such as contrast sensitivity [23] show that there are changes to visual perception related to malnutrition, indicating that the visual mechanisms of this afflicted population may be related to neurophysiological changes in visual cortical areas, such as the primary visual cortex [13].

From the research hypothesis, it was expected that the nutritional deficit during the critical period of the visual system development might affect eye movement. Such results indicate that PEM may promote structural and functional changes to the central nervous system [1, 13]. Visual processing depends on the functioning of the retinae, optical tracts and visual cortex [24, 25]. Then, impairments in every affection to these structures (detachment, degenerations, inflammations, scars of the central part of the retinae, optical neuritis or compromising of the axons related to the ganglion cells of the fovea, lesions affecting the visual cortex or other parts, etc.) or when the development of neuronal competencies itself is imperfectly fulfilled.

Activities such as the maze test and the seven errors test demand mastery of the executive function, one of the most complex aspects of cognition, encompassing information selection, integration of current information to those previously memorised, planning, monitoring and cognitive flexibility [26]. As such, although not being able to extrapolate results, we may indicate the possibility of the existence of deficits in “superior” processing cortical regions, as well as in the “lower” level processing [27, 28].

Supervision in the school with the participant children and collaborating teachers allow us to understand that the abilities of visual perception include capacity to recognise images once seen without attaching meaning to them and recognizing the same symbol in different forms. Furthermore, students with visual perception impairment normally have difficulty with visual memory and visualisation. One of the most serious consequences of malnutrition is the increase in the risk of school failure. A delay in the functional development of the nervous system is critical for the future and success of the affected child.

The growth of a child may be evaluated by the anthropometric examination, which shows eventual deficits in weight and/or height and, therefore, serve for presumable malnutrition processes to be identified. Current malnutrition is when children do not present the ideal weight for their height. Prior malnutrition is defined as a condition in which a child is small for its age and genetic lineage, but does not present specific symptoms and clinical signs, in addition to delays in growth and chronic malnutrition experienced in the past and continuing into the present [13]. Some students with impairment in their visual function also have problem with spatial relations. It is difficult for them to handle concepts as size, form and distance.

As the development of the central nervous system may be affected by nutritional jeopardy of the child in an early post-natal period, it is believed that the visual system may be impaired in its development and develop with ocular sequelae [5].

The results presented show impairments to the visual system, specifically in the eye movement of children with previous PEM for the three test types, supporting the findings. However, it is important that further and more thorough studies be made, with a larger sample number, changes to pupil and adding of other parameters of eye movement, such as length and saccadic speed. These results should be interpreted cautiously.

From this study, we have direct application in the fields of neuroscience, sensorial psychology, public health, among other areas, in addition to allowing for improving specific tests for sensorial disturbances. The study shows us the paucity of information that reaches teachers, so we believe this research has great relevance concerning the cognitive development of children, since this problem affects children’s school organisation and constitution entirely.

## Acknowledgements

National Scientific and Technological Development Council (CNPq), Brazil (432891/2018-8 and 305258/2019-2).

## Conflicts of interests

None.

## Notes

### Competing Interest Statement

The authors have declared no competing interest.

